# Enhancing the protein fitness of interferon-lambda through computational design and glyco-engineering for prophylactic nasal drugs against respiratory viruses

**DOI:** 10.1101/2025.04.16.649247

**Authors:** Jeongwon Yun, Seungju Yang, Jae Hyuk Kwon, Luiz Felipe Vecchietti, Meeyoung Cha, Hyun Jung Chung, Ji Eun Oh, Ho Min Kim

## Abstract

Interferon-λ (IFN-λ), a type III interferon that selectively targets epithelial cells, holds strong potential as an intranasal antiviral due to its ability to suppress respiratory virus replication without inducing systemic inflammation. However, clinical translation of human IFN-λ3 (hIFN-λ3) is hindered by limited thermostability, protease susceptibility, and rapid mucosal clearance. In this study, instability-prone elements in hIFN-λ3 are eliminated through artificial intelligence (AI)-based backbone remodeling and targeted surface hydrophobic patch engineering. A protease-sensitive loop is replaced with a de novo α-helix, which shields neighboring hydrophobic patches and forms a new hydrophobic core, yielding an engineered variant (hIFN-λ3-DE1) with enhanced thermostability (Tm >90 °C), protease resistance, and preserved antiviral activity and structural integrity even after extended heat stress (two weeks at 50°C). Further glyco-engineering introduces an N-linked glycan at a site distant from receptor-binding interfaces, improving solubility, production yield, and diffusion through synthetic nasal mucus. Intranasal administration of the resulting variant (G-hIFN-λ3-DE1) enables effective mucosal penetration and provides robust prophylactic protection against influenza A virus infection in an in vivo mouse model. These findings highlight a robust and versatile strategy that combines AI-driven structural design with glyco-engineering to develop scalable, bioavailable, and functionally enhanced nasal biologics for respiratory virus prophylaxis.

## 1. Introduction

Viral infections pose a persistent and evolving threat to global health, demanding innovative strategies for broadly applicable antiviral therapeutic and prophylactics. Infected viruses are recognized by pattern recognition receptors, including Toll-like receptors (TLRs) and RIG-I-like receptors (RLRs), which trigger the expression of interferons (IFNs)—key cytokines that induce antiviral immunity[1,2]. Among the three classes of IFNs (types I, II, and III), type I IFNs (IFN-α and IFN-β) are well-characterized for their systemic antiviral defense mediated via a ubiquitously expressed receptor complex (IFNAR1 and IFNAR2)[3]. In contrast, type III IFNs (IFN-λs) signal through a receptor complex (IFN-λR1 and IL-10Rβ)[4,5] (Figure 1a) predominantly expressed on mucosal surfaces, such as the respiratory and gastrointestinal epithelia, allowing IFN-λs to induce localized, but potent antiviral responses with minimal systemic inflammation[6–8].

**Figure 1.**
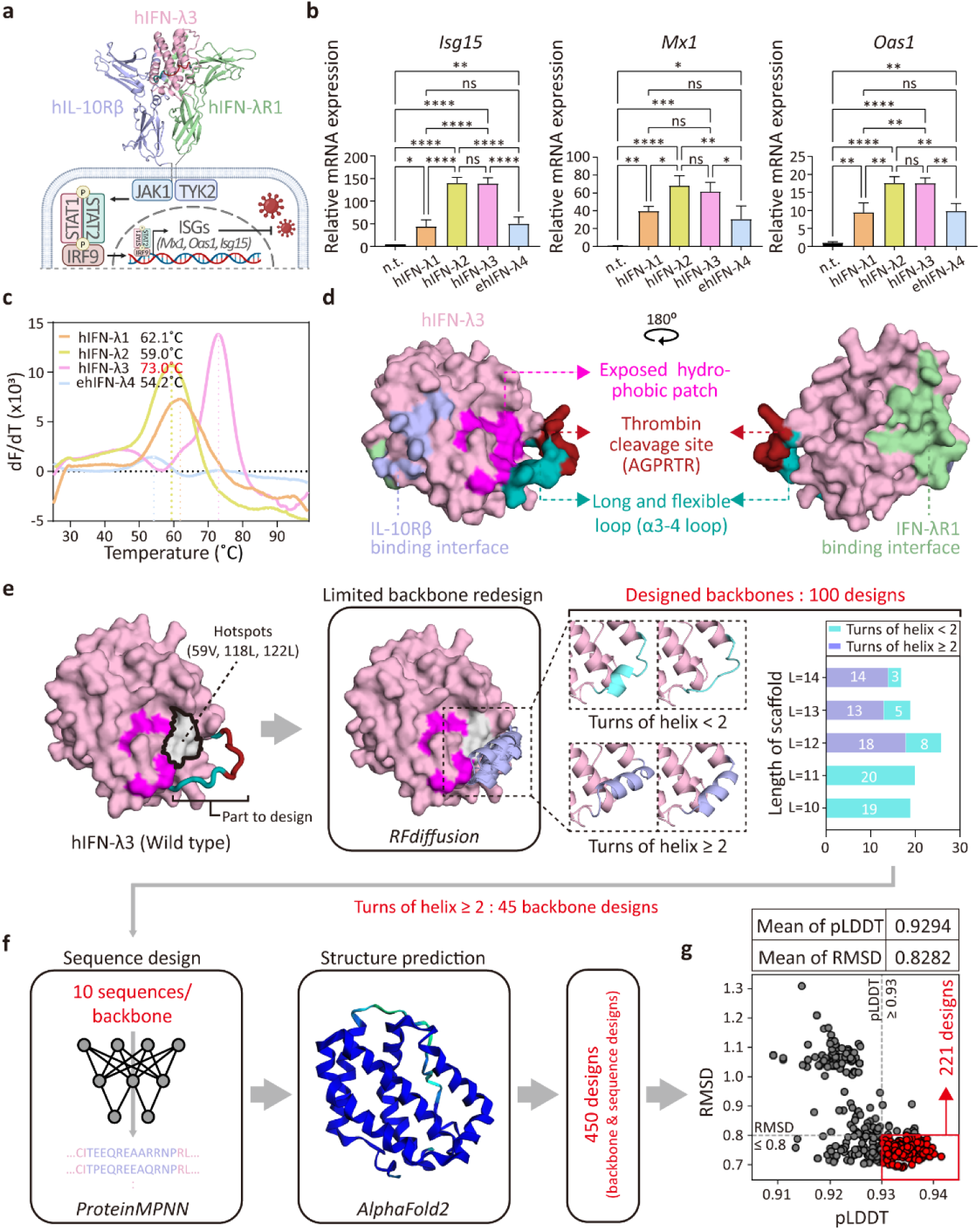
Structure analysis and computational design strategy for thermostable and proteolysis-resistant hIFN-λ3. **a** The 3D structure of the hIFN-λ3/hIFN-λR1/hIL-10Rβ (PDB code: 5T5W)[69] complex and its downstream signaling pathway. **b** Induction of representative ISG expression (*Isg15, Mx1, and Oas1*) in HNEpCs after 12-h treatment with recombinant hIFN-λs WT, analyzed by RT-qPCR (*n* = 3). mRNA levels were calculated relative to non-treated controls and normalized to human *18s rRNA* expression. **c** First-derivative plots (dF/dT) of thermal shift assays for calculating melting temperatures (Tm). 12.5 μg of each hIFN-λ was mixed with 2.5 μL of diluted Protein Thermal Shift™ Dye. Tm values, corresponding to the peak dF/dT temperature, are indicated. **d** Surface representation of hIFN-λ3 (PDB code: 3HHC)[65]. The structure of the α3-4 loop, which is missing in the crystal structure, is predicted by AlphaFold2 (AF2). The binding sites of hIFN-λ3 for IL-10Rβ and IFN-λR1 are highlighted in purple and lime, respectively. The exposed hydrophobic patch (magenta), thrombin cleavage site (red), and flexible α3–4 loop (cyan) are shown. **e** Targeted backbone redesign using RFdiffusion. The designated hotspot residues (59V, 118L, and 122L) on the exposed hydrophobic patch for RFdiffusion are highlighted in light gray. The resulting 100 designed backbones are categorized by scaffold length and the number of α-helix turns at the redesigned backbone. **f** Sequence design of 45 redesigned backbones (10 sequences/1 redesigned backbone) using ProteinMPNN, followed by structure prediction with AlphaFold2 (AF2). **g** Scatter plot showing AlphaFold2-predicted confidence (pLDDT) versus RMSD (AF2 prediction compared to the generated backbone) for 450 designed hIFN-λ3 variants. Dashed lines indicate selection thresholds (pLDDT ≥ 93, RMSD ≤ 0.8), with 221 selected designs in the lower-right quadrant. All data represent mean ± SD from independent experiments. Statistical analysis was performed using one-way ANOVA followed by Tukey’s multiple comparisons test (0.01<\**P*<0.1, 0.001<\*\**P*<0.01, 0.0001<\*\*\**P*< 0.001, \*\*\*\**P*<0.0001 vs. control; ns, not significant).

Emerging evidence underscores the central role of IFN-λs in respiratory antiviral defense, particularly against high-burden pathogens such as influenza A virus (IAV), SARS-CoV-2, and MERS-CoV[9–11]. Clinical studies have demonstrated that elevated levels of IFN-λ correlate with reduced viral loads, enhanced viral clearance, and improved clinical outcomes in COVID-19 patients[12–14], proposing its frontline role in suppressing viral spread within the respiratory epithelium[15,16]. These findings highlight IFN-λ as a promising candidate for broad-spectrum antiviral therapeutics targeting the respiratory epithelium. However, clinical trials of subcutaneous pegylated IFN-λ1 have yielded mixed results[17–19], attributed to the poor delivery to the primary sites of viral infection and replication—the nasal and upper respiratory epithelium.

Nasal delivery, a non-invasive and efficient route to the site of respiratory virus infection and replication, offers the potential for IFN-λ-based therapies[20,21]. Unlike systemic administration, intranasal delivery enables direct access to epithelial IFN-λ receptors, ensuring rapid antiviral action while minimizing systemic exposure. Recent studies have highlighted the efficacy of intranasally delivered recombinant IFN-λ2 in reducing viral loads and mitigating lung damage caused by SARS-CoV-2 variants[22]. Despite this promise, the development of nasal protein therapeutics encounters significant challenges, including protein instability, degradation by nasal proteases, and mucociliary clearance[23–25].

In this study, we employed AI-based protein design and glyco-engineering to overcome these barriers and enhance the therapeutic fitness of human IFN-λ. Our engineered IFN-λ exhibits remarkable stability, with a melting temperature (Tm) exceeding 90°C and resistance to proteolytic degradation, retaining full biological activity even after two weeks incubation at 50°C. Additionally, the engineered IFN-λ demonstrates superior production yields and improved diffusion efficiency across the mucosal barrier. Intranasal administration of this engineered IFN-λ provided robust protection against IAV infection in mouse models, significantly reducing viral burden and associated pathology. These findings establish a new paradigm for the development of protein-based nasal therapeutics, offering a scalable, stable, and effective prophylactic solution against respiratory viruses, including emerging variants. Moreover, our work underscores the transformative potential of computational protein design in addressing critical challenges in antiviral drug development, facilitating the way for next-generation mucosal immunotherapies.

## 2. Results

### 2.1. Comparative analysis identifies hIFN-λ3 as an optimal antiviral candidate for enhancing protein fitness

Humans possess four functional IFN-λ genes (IFN-λ1 to IFN-λ4), each encoding a cytokine with varying antiviral potency and biophysical characteristics. To identify the most suitable candidate for developing prophylactic intranasal biologics against respiratory viruses, we conducted a comparative analysis of all four human IFN-λ proteins. Recombinant hIFN-λ proteins (human IFN-λ1, 2, 3, and 4) were produced in mammalian cells and purified via affinity and size-exclusion chromatography (SEC) (Figure S1a, Supporting Information). For hIFN-λ4, we expressed a glycoengineered version (eIFN-λ4) as previously described[26], since the wild-type hIFN-λ4 is rarely expressed. SDS-PAGE and SEC analysis revealed molecular weights ranging from 20 to 30 kDa, with hIFN-λ1 and eIFN-λ4 being ∼5–10kDa larger than hIFN-λ2 and hIFN-λ3 due to differential glycosylation (Figure S1a, Supporting Information).

To assess functional activity, we treated primary human nasal epithelial cells (HNEpCs) with each recombinant cytokine and measured STAT1 phosphorylation and induction of key interferon-stimulated genes (ISGs: *Isg15, Mx1, and Oas1*). All four IFN-λ proteins activated STAT1 phosphorylation to a similar extent; however, hIFN-λ2 and hIFN-λ3 induced stronger ISG responses, indicating superior antiviral signaling (Figure 1b and Figure S1b, Supporting Information). In parallel, we evaluated thermal stability by measuring melting temperature (Tm) with Thermal shift assays. Among the four proteins, hIFN-λ3 exhibited the highest thermal stability, with a Tm of 73.0°C, which is substantially higher than hIFN-λ1 (62.1°C), hIFN-λ2 (59.0°C), and eIFN-λ4 (54.2°C) (Figure 1c and Figure S1c, Supporting Information). Based on its robust ISG induction and superior thermal stability, hIFN-λ3 was selected as the lead antiviral candidate for structure-guided computational design aimed at further enhancing protein fitness for nasal delivery.

### 2.2. Structure-guided computational design of thermostable and proteolysis-resistant hIFN-λ3

To elucidate structural features relevant to stability and guide rational design, we analyzed the crystal and AlphaFold2 (AF2)[27]-predicted structures of human IFN-λ1, 2, and 3. These proteins share a conserved fold consisting of five α-helices and two 3_10_ helices connected by flexible loops. The N-terminal helix (α1) and the C-terminal helix (α5 of hIFN-λ1, 2, and 3 and α4 of hIFNL4) interact with IFN-λR1, whereas the α1-α3 helices interact with IL-10Rβ (Figure S2, and S3, Supporting Information). Of note, a flexible loop connecting α3 and α4 helices (the α3-4 loop) contains a thrombin cleavage sequence[28]. Adjacent to this flexible α3-4 loop, several conserved hydrophobic residues (eg., V59, A63, A66, L97, A102, I104, L118, and L122 in hIFN-λ3) on helices α2–α4 form a solvent-exposed hydrophobic patch (Figure 1d, Figure S2 and S3, Supporting Information), which does not participate in receptor binding. Solvent-exposed hydrophobic patches are known to negatively affect protein thermodynamic stability[29–32]. Therefore, we hypothesized that masking this patch through loop redesign could improve overall stability and proteolytic resistance.

To address this, we sought to replace the flexible α3-4 loop of hIFN-λ3 (105Q-116R) with a stable α-helix that eliminates the thrombin cleavage sequence and buries the hydrophobic patch through internal core interactions. Using RFdiffusion[33], we performed backbone redesigns at this loop position with lengths ranging from 10 to 14 residues. Three hydrophobic residues at the center of the patch (V59, L118, L122) were designated as design hotspots to guide packing interactions for shielding the hydrophobic patch. A total of 100 backbone models at the α3-4 loop position were generated across four diffusion time steps (T = 50, 100, 150, and 200) (Figure 1e). Secondary structure analysis with the DSSP algorithm[34,35] revealed that loops of 12–14 residues were most effective at forming stable α-helices. From this subset, 45 backbones containing rigid α-helices with ≥8 helical residues were selected for further sequence design using ProteinMPNN[36], generating 10 unique sequences per backbone while keeping the remaining hIFN-λ3 sequence fixed. The structures of 450 designs were predicted by AF2[27] (Figure 1f), resulting in an average predicted local distance difference test (pLDDT) value of 0.9294 and root mean square deviation (RMSD) of 0.8282. To ensure high confidence and structural fidelity, we applied these average values as filtering metrics (i.e., pLDDT ≥0.93 and RMSD ≤0.8) for further analysis, narrowing the pool to 221 designs (Figure 1g).

To assess whether the redesigned α-helix (termed DE-α4) effectively shielded the hydrophobic patch, we calculated the relative solvent accessibility (RSA) of the hotspot residues[37,38] (Figure 2a). Although no strict RSA threshold exists, we reasoned that reducing solvent exposure of hydrophobic regions would improve folding stability. Designs with average RSA < 0.075 across the hotspot residues were prioritized (Figure 2b). Additionally, to prevent unintended surface exposure of new hydrophobic residues introduced by DE-α4, RSA values of hydrophobic residues in the redesigned helix were evaluated. Lastly, we calculated a total free energy using the Rosetta energy function^[39.40]^ after structural refinement through Rosetta relaxation[41,42]. Interestingly, there was a weak correlation between RSA of hydrophobic residues on DE-α4 and total free energy (Figure 2c). Based on these integrated criteria (RSA of hotspots ≤ 0.1 and Rosetta energy ≤ –477), we selected the five most promising candidates (hIFN-λ3-DE1 to DE5; Figure 2c, Table 1, and Table S2, S3, Supporting Information).

**Figure 2.**
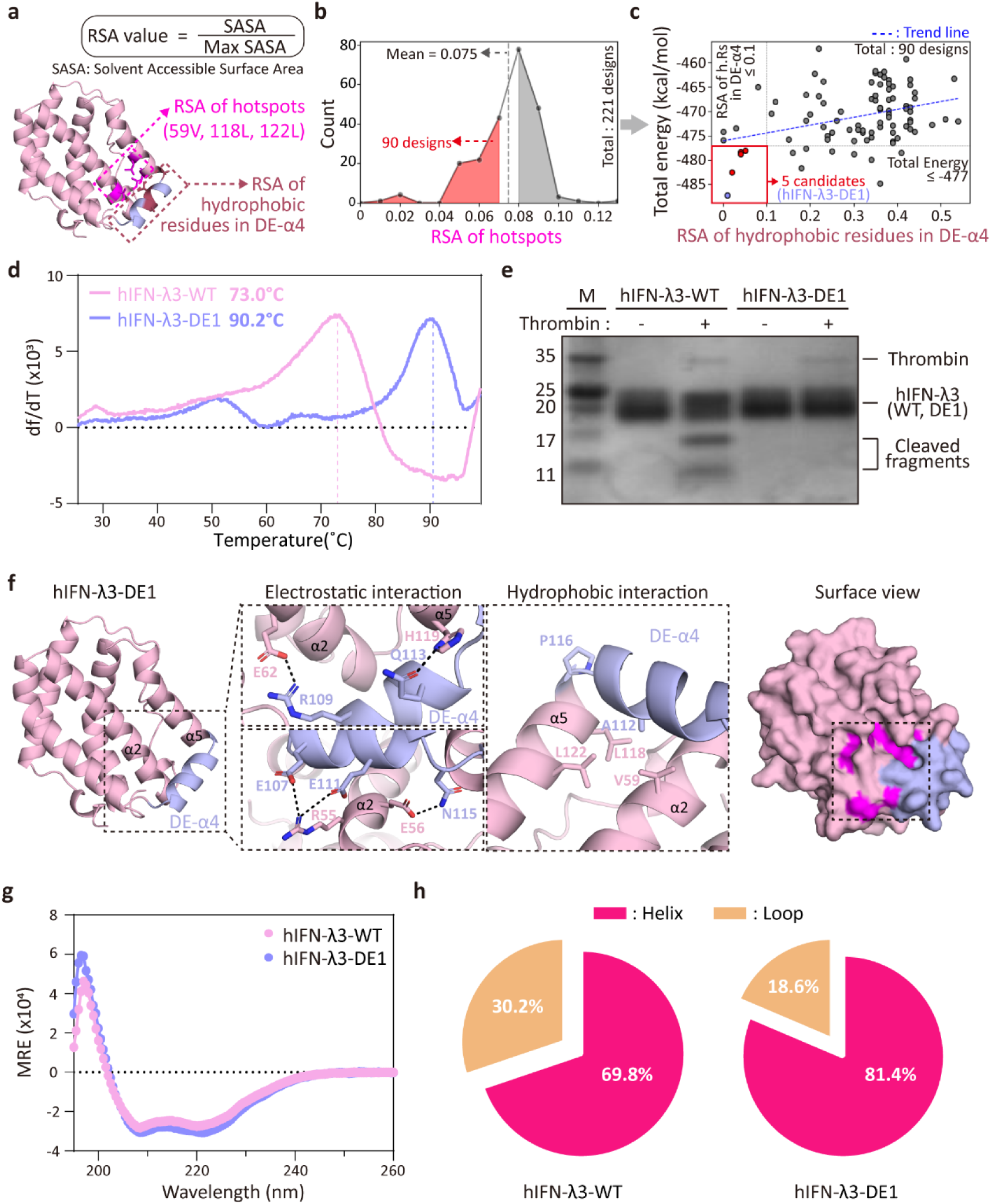
Selection strategy for hIFN-λ3 designs, followed by evaluation of their increased thermal stability, proteolytic resistance, and structural characterization. **a** Schematic showing the calculation of relative solvent accessibility (RSA), defined as the ratio of solvent-accessible surface area (SASA) to the maximum possible SASA (Max SASA) at the designated area. The regions used for RSA calculation of hotspot residues (V59, L118, and L122) and hydrophobic residues in DE-α4 are indicated in magenta and brown, respectively. **b** Histogram showing the distribution of RSA values for the hotspot residues among 221 designed variants. The average “RSA of hotspots” was 0.075, 90 designs below this threshold are highlighted in red. **c** Scatter plot of values for “RSA of hydrophobic residues in DE-α4” vs “total energy (kcal/mol)”. The regression line (y = 16.122x – 476.08, R² = 0.1251) is shown. Five designs satisfying both selection criteria (RSA ≤ 0.1 and total energy ≤ –477) are indicated in red. **d** First derivative (dF/dT) curves of thermal shift assays for determining melting temperature (Tm). The Tm values of hIFN-λ3-WT and hIFN-λ3-DE1 determined are indicated at peak positions. **e** Thrombin susceptibility assay. hIFN-λ3-WT and hIFN-λ3-DE1 were incubated with thrombin (1% [v/v] in PBS) at 37°C for 1 h and analyzed by SDS-PAGE followed by Coomassie blue staining. **f** AF2-predicted model of hIFN-λ3-DE1. Insets highlight stabilizing electrostatic (left) and hydrophobic (middle) interactions between the DE-α4 helix (purple) and neighboring helices (pink). Surface view shows the buried hydrophobic patch (magenta) shielded by DE-α4 (purple). **g** Circular dichroism (CD) spectra of hIFN-λ3-WT and hIFN-λ3-DE1 at 25°C across 195–260 nm. **h** Pie chart showing the secondary structure content of hIFN-λ3-WT and hIFN-λ3-DE1 calculated from CD analysis.

**Table 1:** Comparison of filtering metrics for wild-type and designed hIFN-λ3 candidates

| hINF-λ3<br>(DE-#) | RSA of<br>Hotspots | RSA of Hydrophobic<br>Residues in DE-α4 | Total Energy<br>(kcal/mol ) | ΔE<br>(kcal/mol) |
| --- | --- | --- | --- | --- |
| Wild type | 0.23 | - | -423.434 | 0 |
| **DE-1** | **0.06** | **0.01** | **-487.285** | **-63.851** |
| DE-2 | 0.07 | 0.05 | -478.023 | -54.589 |
| DE-3 | 0.06 | 0.04 | -478.305 | -54.871 |
| DE-4 | 0.07 | 0.04 | -478.717 | -55.823 |
| DE-5 | 0.07 | 0.02 | -482.502 | -59.068 |

ΔE = 'Total Energy' of wild type - 'Total Energy' of hINF-λ3-DE

Synthetic genes encoding five designed hIFN-λ3s were expressed in a mammalian cell and purified (Figure S4a, Supporting Information). All five designs exhibited improved thermal stability compared to wild-type hIFN-λ3 (WT), as assessed by thermal shift assays. The most pronounced enhancement was observed for hIFN-λ3-DE1, which showed a melting temperature (Tm) of 90.2°C—substantially higher than that of hIFN-λ3-WT (Tm = 73.0°C) (Figure 2d and Figure S4b, Supporting Information). This result indicates effective structural stabilization via the DE-α4 design. Furthermore, thrombin cleavage analysis revealed that hIFN-λ3-WT was cleaved into two fragments, whereas hIFN-λ3-DE1 remained intact, demonstrating resistance to proteolysis (Figure 2e). This resistance is attributed to the removal of the thrombin cleavage motif in the α3-4 loop in hIFN-λ3-DE1 and its structural replacement with a well-packed α-helix. Collectively, these results demonstrate that structure-guided computational design can substantially improve the stability and protease resistance of hIFN-λ3. The enhanced biophysical properties of hIFN-λ3-DE1 support its suitability for protease-rich mucosal environments, underscoring its potential as a robust intranasal biologic.

### 2.3. Structural insights into designed hIFN-λ3

To elucidate the molecular basis of the enhanced thermal stability observed in hIFN-λ3-DE1, we examined the AF2-predicted structures of the designed hIFN-λ3s (Figure 2f and Figure S4c, Supporting Information). The hIFN-λ3-DE1 structure revealed newly formed intramolecular interactions between the designed DE-α4 and adjacent helices (α2 and α5), contributing to structural stabilization (Figure 2f). Notably, a sharp kink at P116 in DE-α4, along with multiple electrostatic interactions (E62^α2^-R109^DE-α4^, R55^α2^-E107^DE-α4^ and E111^DE-α4^, H119^α5^-Q113^DE-α4^, and E56^α2^-N115^DE-α4^) locked DE-α4 into a fixed orientation. In addition, A112^DE-α4^ formed hydrophobic contacts with three hotspot residues (V59, L118, and L122), thereby effectively burying the hydrophobic patch within the protein core.

We next validated the structural formation of the designed helical element in hIFN-λ3-DE1 using circular dichroism (CD) spectroscopy. Secondary structure analysis showed a marked increase in helical content—from 69.8% in hIFN-λ3-WT to 81.4% in hIFN-λ3-DE1 (Figure 2g and 2h), supporting the successful loop-to-helix transformation introduced by the DE-α4 design of hIFN-λ3.

### 2.4. Designed hIFN-λ3 retained biological activity under heat stress and long-term storage

Next, we examined whether hIFN-λ3-DE1 retains its biological activity in human nasal epithelial cells (HNEpCs). STAT1 phosphorylation levels were comparable between hIFN-λ3-WT and hIFN-λ3-DE1 (Figure S5a, Supporting Information), suggesting that designed DE-α4 of hIFN-λ3-DE1 preserved the structural integrity of critical functional domains, including receptor-binding interfaces, thereby maintaining full biological activity.

We further assessed resistance to acute heat stress by incubating the proteins at 70°C for 5 minutes prior to HNEpC treatment. Notably, hIFN-λ3-WT lost all ISG-inducing activity, whereas hIFN-λ3-DE1 retained full activity, equivalent to unheated controls (Figure 3a). Furthermore, aggregation analysis revealed that hIFN-λ3-WT aggregated by ∼40% at 60°C and >90% at 70°C within 5 minutes. In contrast, hIFN-λ3-DE1 exhibited remarkable resistance to thermal aggregation, maintaining solubility even at 70°C, with minimal aggregation only at 80°C (Figure 3b and Figure S5b, Supporting Information).

**Figure 3.**
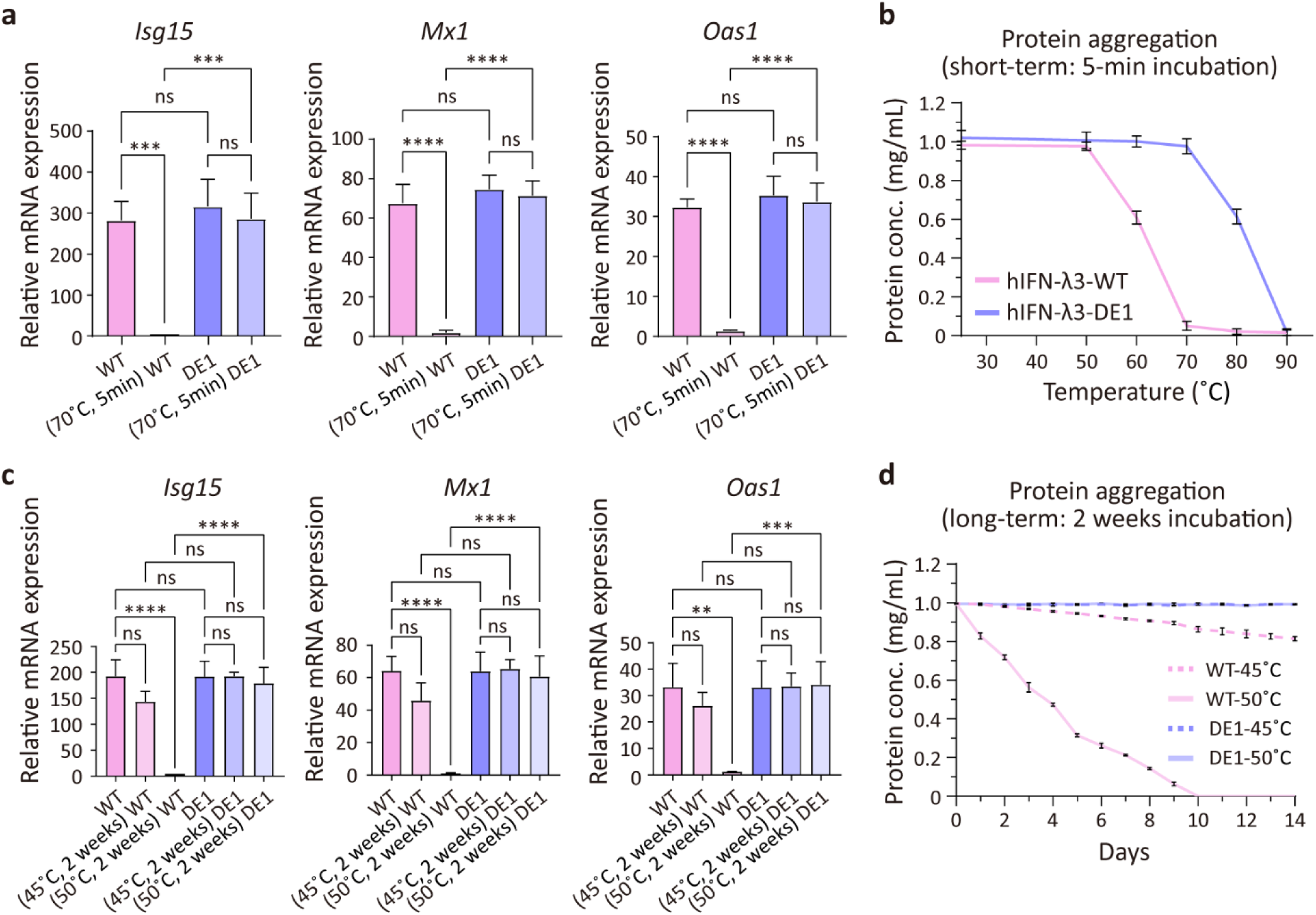
Biological activity and thermal aggregation resistance of hIFN-λ3-DE1 under acute and long-term heat stress. **a** Relative mRNA expression of representative ISGs (*Isg15, Mx1, and Oas1*) in HNEpCs following 12-hour treatment with hIFN-λ3-WT or hIFN-λ3-DE1 (100 ng/mL), with or without short-term heat stress (70°C for 5 minutes). mRNA levels were analyzed by RT-qPCR (*n* = 3), normalized to *18s rRNA*, and expressed relative to non-treated controls. **b** Short-term thermal aggregation profiles of hIFN-λ3-WT and hIFN-λ3-DE1 after 5-minute incubation at the indicated temperatures (25, 50, 60, 70, 80, or 90°C). Residual soluble protein concentrations were quantified (*n* = 3). **c** Relative ISG expression (*Isg15, Mx1, and Oas1*) in HNEpCs treated with hIFN-λ3-WT or hIFN-λ3-DE1 (100 ng/mL) after long-term incubation at 45°C or 50°C for 2 weeks. RT-qPCR was performed as in (a) (*n* = 3). **d** Long-term thermal aggregation of hIFN-λ3-WT and hIFN-λ3-DE1 during 2-week incubation at 45°C or 50°C. Protein solubility was monitored over time (n = 3). All data represent mean ± SD from independent experiments. Statistical analysis was performed by one-way ANOVA followed by Sidak’s multiple comparisons test (0.001<\*\**P*<0.01, 0.0001<\*\*\**P* < 0.001, \*\*\*\**P*<0.0001 vs. control and ns is not significant). n.t., non-treat; WT, hIFN-λ3-WT; DE1, hIFN-λ3-DE1.

To evaluate long-term thermal stability relevant to nasal drug development, we incubated both proteins for 2 weeks at elevated temperatures. Both hIFN-λ3-WT and hIFN-λ3-DE1 retained ISG-inducing activity after 2 weeks at 37°C, comparable to their respective unstressed controls (Figure S5c, Supporting Information). However, hIFN-λ3-WT exhibited reduced activity after 2 weeks at 45°C and lost activity entirely at 50°C. In contrast, hIFN-λ3-DE1 maintained full biological activity even after 2 weeks at both 45°C and 50°C (Figure 3c). Aggregation analysis revealed that hIFN-λ3-WT aggregated by ∼20% at 45°C and reached complete aggregation within 10 days at 50°C, while hIFN-λ3-DE1 remained soluble even after 2 weeks at 45°C and 50°C (Figure 3d). Taken together, these results highlight the superior thermostability of hIFN-λ3-DE1, which maintains both structural integrity and biological activity under rigorous storage and heat stress conditions, underscoring its translational potential as a stable and effective intranasal biologic.

### 2.5. Glyco-engineering improves production and mucosal penetration of designed hIFN-λ3

Glyco-engineering has emerged as a powerful strategy to enhance protein therapeutics by improving solubility, folding stability, and bioavailability, while also reducing aggregation and proteolytic susceptibility under physiological conditions[43–46]. PEGylation, a synthetic mimic of glycosylation, has long been utilized to extend protein half-life and improve diffusion through biological barriers, including the mucus[47–50]. Notably, both PEGylation and glycosylation can create hydrophilic, sterically shielding surfaces that reduce muco-adhesion and facilitate rapid mucosal penetration[51–53]. Here, we sought to apply site-specific glyco-engineering to our optimized IFN-λ3 variant (hIFN-λ3-DE1) for enhancing its manufacturability and mucosal delivery properties as an intranasal biologic.

Based on structural alignment within the IFN-λ family (Figure S2, Supporting Information), we identified Asp35 in hIFN-λ3 as a position analogous to Asn46 in hIFN-λ1, a naturally glycosylated site (Figure S1a and S6a, Supporting Information). As this site lies distal from receptor-binding interfaces, we reasoned that introducing N-glycans here would minimize structural or functional disruption. To this end, we engineered D35N and K37S mutations to create an N-X-S glycosylation motif, generating a glycosylated variant termed G-hIFN-λ3-DE1 (Figure 4a). Expression in mammalian cells and subsequent purification confirmed glycan attachment to G-hIFN-λ3-DE1, as evidenced by increased molecular weight and PNGaseF-dependent band shifts (Figure 4b and S6b, Supporting Information). Importantly, G-hIFN-λ3-DE1 maintained high thermostability (Tm = 90.2°C), comparable to the non-glycosylated hIFN-λ3-DE1 (Figure S6c, Supporting Information), indicating that the introduced glycan did not compromise protein folding. Moreover, glyco-engineering markedly improved protein production yield, achieving a 2.5-fold increase relative to wild-type hIFN-λ3 (Figure 4c), likely due to improved protein solubility conferred by the attached glycans.

**Figure 4.**
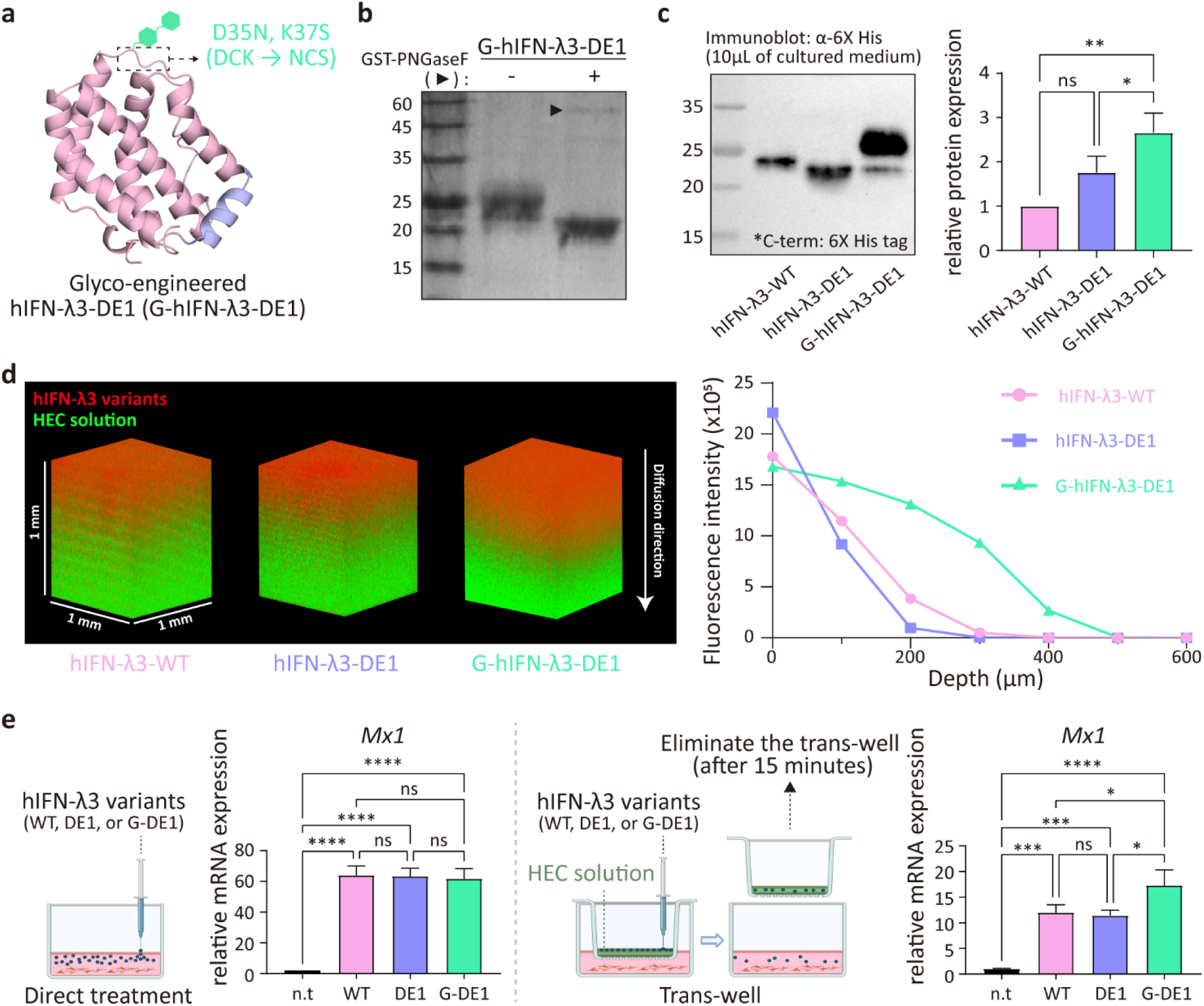
Glyco-engineering enhances production yield and mucosal diffusion of designed hIFN-λ3. **a** Structure-guided glyco-engineering of hIFN-λ3-DE1 to generate the glycosylated variant G-hIFN-λ3-DE1. The mutated amino acid residues for N-glycosylation (D35N/K37S) are indicated. **b** SDS-PAGE analysis of purified G-hIFN-λ3-DE1 before and after PNGaseF-mediated deglycosylation. **c** Expression of hIFN-λ3 variants (WT, DE1, and G-DE1) in Expi293F cells. Same volumes of culture supernatants were collected 4 days post-transfection and analyzed by immunoblotting using an anti-His antibody (*n* = 3). All hIFN-λ3 variants contained a C-terminal 6X-His tag. **d** Confocal imaging of Alexa 647-labeled hIFN-λ3 variants (WT, DE1, and G-DE1) diffusing over 10 minutes through a 1.2% hydroxyethyl cellulose (HEC) matrix pre-labeled with BDP-FL (left). Fluorescence intensity across z-depth is quantified (right). **e** Relative *Mx1* mRNA expression in HNEpCs after 12-hr incubation with hIFN-λ3 variants (100 ng/mL, WT, DE1, and G-DE1). Cells were either treated directly (left) or via a trans-well diffusion system with 500 μm-thick HEC layer (right). In the trans-well setup, protein was applied to the upper insert and removed after 15 minutes and cells were then incubated for 12 hours prior to RT-qPCR (*n* = 3). All data represent mean ± SD from independent experiments. Statistical significance was determined by one-way ANOVA followed by Tukey’s multiple comparisons test (0.001<\*\**P*<0.01, 0.0001<\*\*\**P* < 0.001, \*\*\*\**P*<0.0001 vs. control, and ns is not significant). n.t., non-treat; WT, hIFN-λ3-WT; DE1, hIFN-λ3-DE1; G-DE1, G-hIFN-λ3-DE1.

To evaluate the functional impact of glycosylation on mucosal transport, we assessed the diffusion of G-hIFN-λ3-DE1 through an artificial 3D gel system mimicking the nasal mucus composed of 1.2% hydroxyethyl cellulose (HEC), representing the viscous and glycan-rich environment of the native mucus (1.6 Pa/s)[54,55]. Fluorescently labeled hIFN-λ3-WT, hIFN-λ3-DE1, and G-hIFN-λ3-DE1 were applied atop a 1 mm-thick HEC gel in a microplate, and penetration was examined by confocal microscopy after 10 minutes. G-hIFN-λ3-DE1 exhibited markedly greater diffusion depth compared to both wild-type and non-glycosylated variants (Figure 4d), suggesting that the glycol-engineering promoted more efficient diffusion of G-hIFN-λ3-DE1 through the mucus-like barrier.

We further assessed the biological relevance of mucosal diffusion by comparing Mx1 mRNA induction in HNEpCs treated with hIFN-λ3 variants (hIFN-λ3-WT, hIFN-λ3-DE1, G-hIFN-λ3-DE1) directly or via the HEC gel placed in trans-well inserts. Direct treatment of HNEpCs with each variant resulted in comparable levels of Mx1 transcription, confirming that glycosylation did not affect intrinsic bioactivity (Figure 4e, left). In contrast, in the trans-well system with 500 μm-thickness of HEC gels, G-hIFN-λ3-DE1 induced significantly higher Mx1 expression in HNEpCs relative to hIFN-λ3-WT and hIFN-λ3-DE1 (Figure 4e, right). These results demonstrate that site-specific glyco-engineering improves mucosal penetration and functional delivery of IFN-λ3 across mucus-like barriers, offering a promising strategy for nasal protein drug development.

### 2.6. Intranasal delivery of designed hIFN-λ3 confers effective prophylactic protection against IAV infection in mice

We first examined the ability of the hIFN-λ3 variants to induce interferon-stimulated genes (ISGs) in vivo following intranasal administration. Mice treated with purified hIFN-λ3-WT, hIFN-λ3-DE1, or G-hIFN-λ3-DE1 exhibited significant upregulation of *isg15*, *mx1*, and *oas1* transcripts in the nasal epithelium compared to untreated controls (Figure 5a). Consistent with *in vitro* assay with HNEpCs, hIFN-λ3-WT and hIFN-λ3-DE1 induced comparable levels of ISG expression, suggesting that protein stabilization did not alter basal activity. In contrast, G-hIFN-λ3-DE1 elicited slightly higher *isg15* expression than both non-glycosylated variants, along with a modest increase in *mx1* and *oas1* expression. These results support the notion that glyco-engineering enhances access to epithelial receptors beneath the mucus layer, potentially improving mucosal signal transduction and antiviral efficacy.

**Figure 5.**
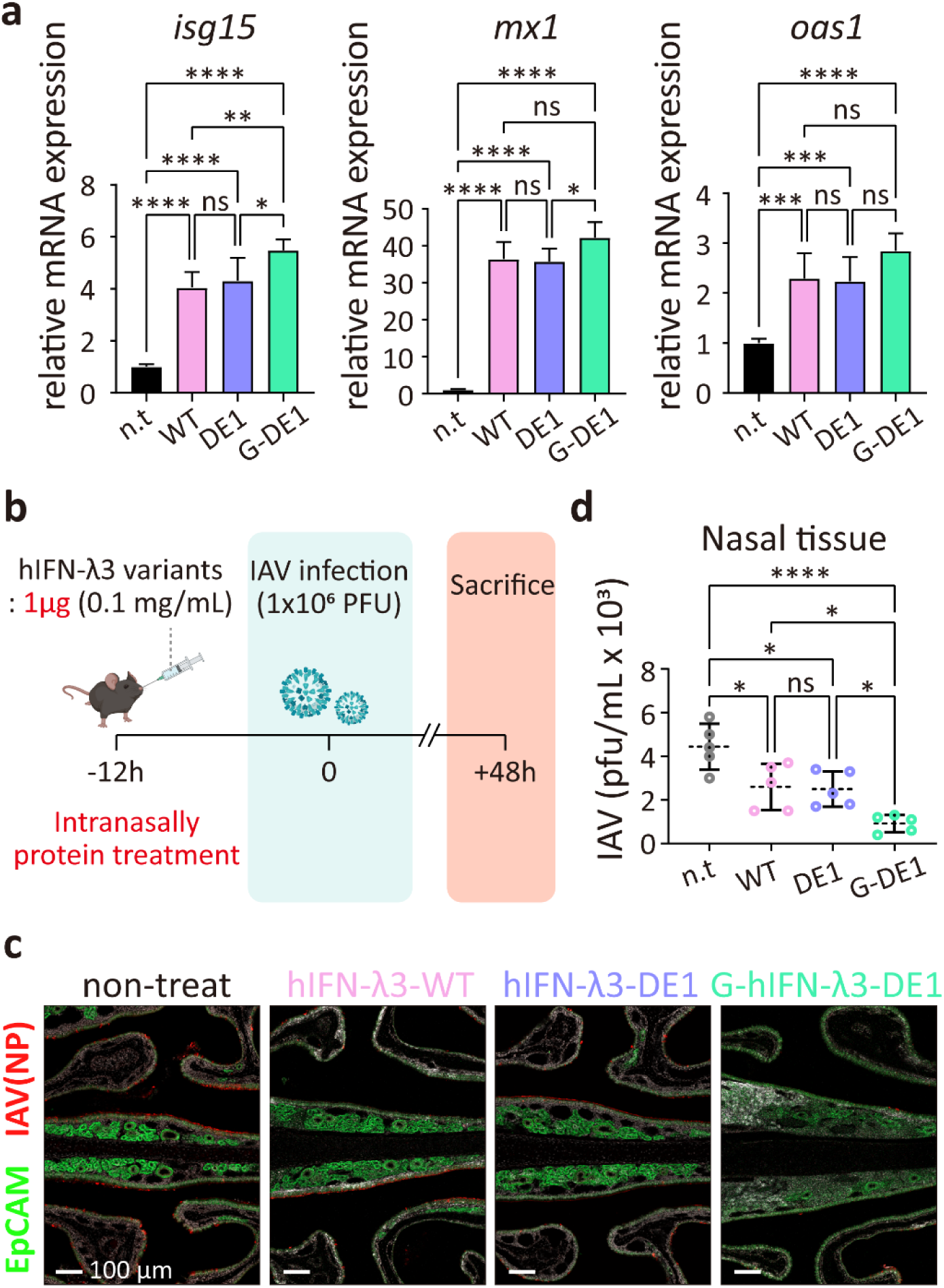
Intranasal delivery of engineered hIFN-λ3 variants provides enhanced prophylactic effects in an IAV-infected mouse model. **a** Relative expression levels of representative ISGs (*isg15, mx1, and oas1*) in mouse nasal turbinates 2 hours after intranasal administration of recombinant hIFN-λ3-WT, hIFN-λ3-DE1, or G-hIFN-λ3-DE1 (5 μg in 10 μL PBS; 0.5 mg/mL). mRNA levels were analyzed by RT-qPCR (*n* = 3), normalized to mouse *gapdh*, and expressed relative to non-treated controls. **b** Schematic experimental design for IAV infection and intranasal administration of hIFN-λ3 variants (WT, DE1, and G-DE1). Mice were intranasally pre-treated with hIFN-λ3 variants (1 μg) 12 hours prior to IAV infection (1×10⁶ PFU), followed by analysis 48 hours post-infection. **c** Immunofluorescence images of nasal tissue sections of IAV-infected mice (dpi=2) intranasally treated with hIFN-λ3 variants (WT, DE1, and G-DE1). Epithelial cells were labeled with anti-EpCAM antibody (green) and influenza A nucleoprotein was detected with anti-IAV antibody (red). Representative images are shown for each treatment group. **d** Quantification of viral titers in nasal tissues at 2 dpi by standard plaque assay (*n* = 5 mice per group). Each dot represents an individual mouse; horizontal lines indicate mean ± SD. All data represent mean ± SD from independent experiments. Statistical significance was determined by one-way ANOVA test followed by Tukey’s multiple comparisons test (0.01<\**P*<0.1, 0.001<\*\**P*<0.01, 0.0001<\*\*\**P* < 0.001, \*\*\*\**P*<0.0001 vs. control and ns is not significant). n.t., non-treat; WT, hIFN-λ3-WT; DE1, hIFN-λ3-DE1; G-DE1, G-hIFN-λ3-DE1

Next, we evaluated the prophylactic effect of three variants in a mouse model of influenza A virus (IAV) infection by assessing viral replication in the nasal epithelium. Mice received 1 μg of hIFN-λ3-WT, hIFN-λ3-DE1, or G-hIFN-λ3-DE1 intranasally 12 hours prior to viral challenge (Figure 5b). At 48 hours post-infection, IAV replication in the nasal cavity was visualized by confocal microscopy using IAV nucleoprotein (NP) staining (Figure 5c). The untreated group showed widespread NP signal (red) throughout the nasal epithelium (green), indicating extensive viral proliferation. However, mice treated with hIFN-λ3-WT or hIFN-λ3-DE1 exhibited moderate reductions in NP signal, while G-hIFN-λ3-DE1 treatment led to a pronounced suppression of viral replication (Figure 5c). To quantitatively confirm these observations, plaque assays were performed using nasal tissue homogenates. Viral titers were significantly reduced in the G-hIFN-λ3-DE1-treated group, whereas moderate reductions were observed in mice treated with hIFN-λ3-WT or hIFN-λ3-DE1 (Figure 5d). Taken together, these findings demonstrate that glyco-engineered hIFN-λ3-DE1 exhibits superior *in vivo* prophylactic protection against IAV infection when administered intranasally. This enhanced efficacy is likely due to improved diffusion through the viscous, glycan-rich nasal mucus layer, enabling more efficient access to epithelial receptors and thereby enhancing antiviral signaling. These properties highlight the therapeutic potential of G-hIFN-λ3-DE1 as a stable and effective prophylactic biologic for respiratory virus infections.

## 3. Discussion

In this study, we applied AI-based protein design and glyco-engineering to enhance the fitness of human interferon-λ3 (hIFN-λ3). The engineered hIFN-λ3 variant combines enhanced biophysical properties, efficient mucosal diffusion, and superior *in vivo* antiviral efficacy, supporting its potential as an intranasal prophylactic against respiratory viruses.

A major advance of this work is the integration of AI-based backbone reconfiguration with targeted surface patch engineering to address long-standing challenges in protein stability. Flexible loops susceptible to proteolysis and solvent-exposed hydrophobic patches are well-known features that compromise protein stability and integrity. However, conventional engineering approaches have been limited in addressing these elements, largely due to the inherent difficulty of precisely modifying backbone conformations in a desired manner. Strategic sequence redesign and backbone remodeling—altering the structural framework of a protein—has been shown to significantly enhance thermostability and protease resistance[56]. In this study, we also employed state-of-the-art AI-based protein design tools, including RFdiffusion[33], ProteinMPNN[36] and AlphaFold2[27], to replace the α3-4 loop in hIFN-λ3 with a de novo α-helix. This loop-to-helix transformation not only eliminated a thrombin cleavage motif, but also created a new, well-packed hydrophobic core by shielding a surface-exposed hydrophobic patch that had rendered the wild-type protein unstable. Additional proline substitutions at formerly flexible sites further rigidified the local structure and reduced conformational entropy. As a result, the engineered hIFN-λ3-DE1 exhibits exceptional thermostability (melting temperature >90°C) and robust resistance to proteolytic degradation, surpassing the stability achievable by conventional point mutations. These enhancements enabled the hIFN-λ3-DE1 to retain structural integrity and full antiviral potency even after prolonged thermal stress (eg. 2 weeks at 50°C), highlighting its suitability for long-term storage at ambient temperature without refrigeration. Our findings demonstrates how AI-based backbone reconfiguration and hydrophobic patch shielding can be synergistically applied to dramatically enhance protein fitness, establishing a broadly applicable framework for designing stable and effective protein therapeutics for harsh biological environments.

Glyco-engineering is a well-established strategy that can simultaneously enhance a protein’s stability, pharmacokinetics, and in vivo efficacy[57]. For example, Fung et al. introduced N-linked glycan sites into an Fc-fusion of GDF15 and observed markedly improved expression titers, solubility, and sialic acid content, resulting in an extended serum half-life compared to the non-glycosylated counterpart[58]. Similarly, glyco-engineered interferon-β variants—with additional N-linked glycosylation—retained full receptor activity while exhibiting enhanced structural stability and pharmacokinetic profiles[59]. A particularly notable success of therapeutic biologic is darbepoetin-α (Aranesp), a hyper-glycosylated erythropoietin analog engineered with two extra N-glycans, which demonstrated a ∼3-fold increase in serum half-life over regular EPO, enabling less frequent dosing[60]. Building on these precedents, glyco-engineering of hIFN-λ3-DE1 similarly provided both functional and practical advantages. Introduction of *de novo* N-linked glycan at a site distant from receptor-binding interfaces improved protein production and solubility, making the G-hIFN-λ3-DE1 variant well-suited for scalable manufacturing. In addition, G-hIFN-λ3-DE1 diffused more efficiently through a nasal mucus-mimicking artificial gel system than its non-glycosylated counterpart—likely due to the favorable physicochemical properties—demonstrating improved mucosal penetration, a key requirement for intranasal prophylaxis against respiratory viral infections. These enhancements translated into superior *in vivo* protection against influenza A virus (IAV) following intranasal administration. This strategy aligns with previous findings that hydrophilic moieties such as PEG or glycans can increase solubility and bioavailability. However, compared to post-expression modifications like PEGylation, the genetically encoded glycan offers a more streamlined approach to tuning pharmacokinetics and mucosal interactions without additional processing steps. Collectively, these results position glyco-engineering as a robust and versatile strategy for developing scalable, bioavailable, and functionally optimized nasal prophylactic biologics, complementing the stability gains achieved through AI-based backbone reconfiguration.

An additional strength of our approach lies in the immunological properties of IFN-λ3, which have been increasingly recognized as highly favorable for mucosal antiviral defense. Unlike type I interferons, which can induce broad systemic immune activation, type III interferons act primarily on epithelial cells due to the restricted expression of the IFN-λ receptor complex (IFNLR1/IL-10Rβ), thereby confining their activity to barrier tissues such as the respiratory tract[61,62]. This selective action enables potent antiviral signaling with minimal off-target inflammation. In support of this, intranasally administered IFN-λ2 has been shown to confer robust protection against multiple SARS-CoV-2 variants, including Omicron, in murine models[22]. Notably, such protection was achieved without inducing pro-inflammatory cytokines or damaging lung tissue, underscoring the tissue-protective and inflammation-sparing nature of IFN-λ responses. In these models, IFN-λ acted predominantly on radio-resistant, non-hematopoietic epithelial cells and accelerated viral clearance while preserving epithelial integrity[6,63]. Moreover, systemic administration of pegylated IFN-λ1 in clinical trials has demonstrated antiviral efficacy comparable to type I IFNs, but with significantly fewer adverse effects[17], further supporting the safety profile of this cytokine class. Beyond its direct antiviral effects, IFN-λ plays a crucial role in enhancing adaptive immunity. Mice lacking IFN-λ signaling showed impaired CD8^+^ T cell and antibody responses following infection with a live-attenuated influenza virus[64]. Moreover, intranasal administration of IFN-λ alongside subunit vaccines resulted in a 15-fold increase in serum IgG level and mucosal IgA production, markedly improving survival after challenge with virulent influenza strains. Notably, this IgG-boosting effect of IFN-λ was not observed when the vaccine was delivered via intraperitoneal or subcutaneous routes, highlighting the critical role of IFN-λ in potentiating adaptive immunity at mucosal surfaces. Building on these insights, our glyco-engineered hIFN-λ3 variant is designed for intranasal delivery to maximize localized action while minimizing systemic exposure. To reduce the risk of immunogenicity, we limited sequence alterations to a small region—specifically, 12 amino acids for structural stabilization and 2 for glycan addition—and designed the protein to be effective at low doses. While anti-drug antibody responses cannot be entirely ruled out, the minimal modifications and tissue-localized mechanism of action are expected to reduce such risks. Taken together, our findings reinforce the growing consensus that IFN-λ is an ideal candidate for mucosal immunoprophylaxis, offering targeted and durable protection against diverse respiratory virus with minimal inflammatory consequences.

Our engineered hIFN-λ3 shows strong potential as a first-in-class intranasal prophylactic agent against diverse respiratory virus. The next steps toward clinical translation will involve rigorous preclinical safety evaluations and proof-of-concept efficacy studies in humans to confirm its protective effects and tolerability. Successfully developing this biologic could introduce a new approach to respiratory virus prophylaxis by providing a fast-acting, broadly protective shield at the site of infection, thereby strengthening our defenses against future respiratory pandemics.

## 4. Experimental Section/Methods

### Computational design process

RFdiffusion[33] was employed to design new scaffolds to replace the flexible loop in hIFN-λ3, which contains the thrombin cleavage consensus sequence, with a stable α-helix for concealing the nearby exposed hydrophobic patch. In brief, the B chain of the crystal structure of hIFN-λ3 (PDB code: 3HHC)[65] was used as the input structure. We tasked RFdiffusion with building 10-14 residues to replace the loop connecting α3 and α4 (the α3-4 loop, residues 105-116, L=12) (Figure S3, Supporting Information). Three hydrophobic residues (Val59, Leu118, and Leu122), located at the center of the solvent-exposed hydrophobic patch, were designated as spatial hotspots to guide backbone generation by RFdiffusion. Twenty-five backbones replacing the α3-4 loop of hIFN-λ3 were generated at each diffusion step (T=50, 100, 150, and 200) with the final contig being ‘B4-104/10-14/B117-163’, resulting in a total of 100 designed hIFN-λ3s. Next, we selected designed hIFN-λ3 candidates containing a rigid α-helix with more than two turns (> 8 amino acids) at the initial α3-4 loop position. To do so, we analyzed their secondary structures with the DSSP algorithm[34,35], which identifies repetitive patterns of intra-backbone hydrogen bonds by calculating the electrostatic energy based on the distance between the carbonyl group atoms (C, O) and the amide group atoms (N, H) of the protein backbone. Following the selection of 45 designed hIFN-λ3 candidates, 10 sequences per designed backbone were generated with ProteinMPNN[36], keeping the original sequence of other hIFN-λ3 parts fixed (sampling temperature of 0.1 and backbone noise of 0). Subsequently, their structures were predicted using AlphaFold2[27] with the single sequence option. Structural templating with multiple sequence alignments (MSAs) was not used for structure prediction. A total of 450 designs with backbone and sequence redesign at the α3-4 loop of hIFN-λ3 were filtered to Cα RMSD ≤ 0.8 and pLDDT ≥ 93.0, resulting in 221 designed hIFN-λ3 candidates.

To further select the final candidates with lower energy by minimizing the exposure of hydrophobic amino acids, we used the relative solvent accessibility (RSA) value[37,38], which represents the exposure of amino acids by comparing the solvent-accessible surface area (SASA) at the designated area to their maximum exposure (Max SASA[38]), and the total energy score calculated by energy functions[39,40] in Rosetta[66]. First, we calculated the RSA value for the hotspot residues (Val59, Leu118, and Leu122), the hydrophobic amino acids at the center of the exposed hydrophobic patch. We filtered the 221 designs to those with an RSA of hotspots ≤ 0.075 (average value among the 221 outputs), resulting in 90 designs. Next, we calculated the solvent exposure of the hydrophobic amino acids (Ala, Gly, Val, Ile, Leu, Phe, and Met) on the newly designed α-helix (RSA of hydrophobic residues in DE-α4) among the resulting 90 designs. Lastly, we refined the structures of 90 designs using the relax option[41,42] in Rosetta[66] and calculated the total energy score using the ’ref2015’ (Rosetta energy function)[39,40]. Consequently, 90 designs were filtered with RSA of hydrophobic residues in DE-α4’ ≤ 0.1 and total energy ≤ -477 kcal/mol, resulting in five final designed hIFN-λ3 candidates (hIFN-λ3-DE1, 2, 3, 4, and 5).

### Cell lines and cell culture

HNEpC cells (C-12620, PromoCell) were cultured in Airway Epithelial Cell Growth Medium (C-21060, PromoCell) at 37°C in a humidified 5% CO2 incubator. Expi 293-F cells (A14527, Thermo Fisher Scientific) were maintained in Expi293 Expression Medium (A1435102, Thermo Fisher Scientific) under shaking conditions at 37°C in a humidified 8% CO_2_ incubator.

### Constructs for recombinant hIFN-λs and designed hIFN-λ3 protein expression

Full-length human IFN-λ1, -λ2, and -λ3, along with designed hIFN-λ3 variants, were cloned into the HindIII and NotI sites of a modified pcDNA3.1 vector (#V79020, Invitrogen) incorporating a TEV protease cleavage site and C-terminal 6×His tag. Glycoengineered human IFN-λ4 (ehIFN-λ4: L28N, P73N)[26] was cloned into the HindIII and XbaI sites of the modified pcDNA3.1 vector (#V79020, Invitrogen) containing the thrombin cleavage sequence and Protein A tag.

### Expression and purification of recombinant hIFN-λs and designed hIFN-λ3 candidates

Recombinant plasmids (1 μg) were transfected into 2.5×106 Expi293-F cells (#A14527, Thermo Fisher Scientific) using the ExpiFectamine 293 Transfection kit (#A14524, Thermo Fisher Scientific). Cells were cultured at 37°C and 8% CO_2_ with shaking (orbital shaker, 120 rpm) for 4 days. After centrifugation to remove the cells, the supernatants (hIFN-λ1, 2, and 3 and designed hIFN-λ3 candidates) were loaded onto the Ni-NTA agarose affinity column (#30210, QIAGEN). After washes with 10 column volumes of wash buffer (PBS, 25 mM imidazole), proteins were eluted with elution buffer (PBS, 250 mM imidazole). Eluted proteins were treated with TEV protease (1% [v/v]) at room temperature for 2 hours to remove the C-terminal His tag. For purifying the ehIFN-λ4, the supernatants were loaded onto IgG Sepharose 6 Fast Flow (#17-0969-01, Cytiva). After washes with 10 column volumes of wash buffer (PBS), the Protein A─fused eIFN-λ4─bound resin was incubated with thrombin (1% [v/v] in PBS) at 4°C for 16 h to remove the C-terminal Protein A tag. After incubation, proteins were eluted with an elution buffer (PBS). All of eluted recombinant proteins were further purified by size-exclusion chromatography using a Superose 6 Increase 10/300 GL column (#GE29-0915-96, Cytiva) or Superdex 200 Increase 10/300 GL column (#GE28-9909-44, Cytiva) equilibrated with PBS. The peak fractions were pooled and concentrated to ∼1 mg/mL using an Amicon Ultra centrifugal filter (#UFC8010, Millipore). For deglycosylation analysis, hIFN-λs were incubated with GST-PNGaseF (10 μg/mL) at 37°C for 3 h.

### Immunoblotting of intracellular signal activation in HNEpCs

HNEpC cells (#C-12620, PromoCell) were cultured in basal Airway Epithelial cell Growth Medium (#C-21060, PromoCell) without supplement mix for starvation at 37°C in a humidified 5% CO_2_ incubator for 12 h. After starvation, the cells were incubated with 100 ng/mL of recombinant hIFN-λs for 1 h. Cells were then washed with cold PBS and lysed with lysis buffer (10 mM Tris-Cl pH 7.4, 150 mM NaCl, 5 mM EDTA, 10% glycerol, 1% Triton X-100, protease inhibitor, phosphatase inhibitor). Lysates were mixed with a 5× SDS sample buffer containing 2-mercaptoethanol and heated at 90°C for 5 min. Samples were electrophoresed on a 10% SDS protein gel, and transferred to a 0.45-μm nitrocellulose membrane. The membrane was blocked by incubation with 5% (w/v) skim milk in Tris-buffered saline containing 0.1% Tween-20 at room temperature for 1 h and then incubated with Stat1 (D1K9Y) rabbit monoclonal antibody (mAb; 1:1000 dilution, #14994, Cell Signaling), or phospho-Stat1 (Y701) rabbit mAb (1:1000 dilution, #9167, Cell Signaling), or beta actin (C4) mouse mAb (1:1000 dilution, #sc-47778, Santa Cruz) at 4°C for 8 h. Blots were then incubated with horseradish peroxidase (HRP)-conjugated goat anti-rabbit IgG antibody (1:5000 dilution, #12-348, Sigma-Aldrich) or HRP-conjugated goat anti-mouse IgG antibody (1:5000 dilution, #HAF007, R&D Systems). Immunoreactive Stat1, phosphor-Stat1 and beta actin were visualized using enhanced chemiluminescence. Signal intensities were quantified using ImageJ (v1.54) and GraphPad Prism (v8.0.2) software.

### RT-qPCR of representative ISGs in HNEpCs

Starved HNEpCs (#C-12620, PromoCell) were incubated with 100 ng/mL of recombinant hIFN-λs (wild type, designed and heat-incubated proteins) for 12 h. Total RNA was extracted using Trizol reagent (#15596026, Invitrogen) following the manufacturer’s protocol. Extracted RNA was reverse-transcribed using an oligo-dT primer and SuperScript III First-strand Synthesis System (#18080-051, Invitrogen) according to the kit instructions. Next, qPCR was performed using PowerUP SYBR Green Master Mix (#A25741, Applied Biosystems) on a QuantStudio 5 Real-Time PCR System (Applied Biosystems). The RT-qPCR primers are listed in Table S1 (Supporting Information). Ct values were normalized against *18S rRNA*, and the relative mRNA expression levels compared to untreated samples (PBS) were calculated using the ΔΔCt method (fold expression=2^−ΔΔCt^). Statistical significance for time-course qPCR was determined using one-way ANOVA.

### Thermal shift assay for measuring Tm

Thermal shift assays were conducted using the QuantStudio 5 Real-Time PCR System (Applied Biosystems) and the Protein Thermal Shift Dye Kit (#4461146, Applied Biosystems). Each 20 μL reaction contained 12.5 μL of protein (1 mg/mL), 2.5 μL of diluted Protein Thermal Shift™ Dye (final concentration: 8×), and 5 μL of Protein Thermal Shift™ Buffer. Samples were loaded in a 96-well PCR plate and subjected to a two-step melt curve: step 1, 25°C for 2 min; step 2, temperature increase at 0.05°C/s to 99°C, then held at 99°C for 2 min. The filter was set at ROX with no quencher and no passive filter. Representative fluorescence spectra from ≥3 independent experiments were plotted. Tm values were determined by differentiating melt curves in GraphPad Prism 8.0.

### Thrombin susceptibility test

Purified hIFN-λ3-WT and hIFN-λ3-DE1 were incubated with thrombin (1% [v/v] in PBS) at 37°C for 1 h. Samples then were incubated at 99°C for 5 min with 5× SDS sample buffer containing a 2-mercaptoethanol and electrophoresed on a 4%─20% Mini-PROTEAN® TGX™ Precast Protein Gel (4561094, Bio-Rad).

### CD spectra for analyzing secondary structure

The circular dichroism (CD) spectra of hIFN-λ3-WT and hIFN-λ3-DE1 (1 mg/mL, 60μL) were recorded at 25°C using a Jasco CD J-815 150-L spectropolarimeter (JASCO) in a demountable U-shaped cuvette with a 0.2-mm path length. Measurements were taken from 195 to 260 nm with 1-nm step intervals, and each spectrum was averaged from three scans. Secondary structure contents were analyzed using BeStSel[67] based on the spectra between 195 and 260 nm.

### Protein aggregation test

For short-term incubation, purified hIFN-λ3-WT and hIFN-λ3-DE1 (1 mg/mL, 20μL) were incubated at 50°C, 60°C, 70°C, 80°C, or 90°C for 5 min. For long-term incubation, the purified hIFN-λ3-WT and hIFN-λ3-DE1 (1 mg/mL, 100μL) were incubated at 45°C or 50°C for 2 weeks. The temperature was increased at 0.1°C/s from room temperature (25°C) to the target temperature. After incubation, samples were centrifuged at low speed (3,000 g, 10 min), and the protein concentration in the supernatant was quantified using a DS-11+ Spectrophotometer (DeNovix).

### HEC gel diffusion assay; 3D imaging

For 3D imaging of diffusion in the HEC gel systems, a 1.2% hydroxyethyl cellulose (HEC) solution was prepared by dissolving 0.6 g of HEC powder (#09368, Sigma-Aldrich) in 50 mL of distilled water while heating and stirring at 70°C. For 3D imaging, 150μL of a 1 mM alkyne-BDP-FL solution (#CLK-045, Jena Bioscience) was added into the 1.2% HEC solution, followed by coating the 24-well culture plate (#30024, SPL Life Sciences). For synthesis hIFN-λ3 variants-AF647, hIFN-λ3 variants were reacted with AF647-NHS ester (#A37566, Thermo Fisher) at a molar ratio of 1:4 at room temperature for 2 hours. Unreacted dye was removed using an Amicon Ultra-0.5 centrifugal filter with a 3kDa molecular weight cutoff (#UFC5003, Millipore). Then, 1 μg of AF647-labeled hIFN-λ3 variants was applied to the HEC gels and analyzed by confocal microscopy (LSM 880, Carl Zeiss). Z-stack images were captured, and the fluorescence intensity across various focal planes was quantified using Zen Blue software (Carl Zeiss).

### HEC gel diffusion assay; trans-well model

For the HEC gel diffusion assay in a trans-well system, HNEpC cells (#C-12620, PromoCell) in a 24-well microplate were starved in basal medium for 12 h prior to stimulation. A 500μm-thick layer of 1.2% HEC solution was added to the trans-well inserts (#A37566, SPL Life Sciences) and placed in 24-well microplates. hIFN-λ3 variants (50 ng/5μL) were loaded to the top of the gel in the insert and removed after 15 min incubation. After an additional 12 h incubation at 37°C, total RNA from cells in the lower chamber was extracted using the TaKaRa MiniBEST Universal RNA Extraction Kit (#9767A, TaKaRa). qPCR was performed with PowerUP SYBR Green Master Mix (#A25741, Applied Biosystems) on a QuantStudio 5 Real-Time PCR System (Applied Biosystems). The RT-qPCR primers are listed in Table S1 (Supporting Information). Ct values were normalized to *18S rRNA*, and gene expression was calculated using the ΔΔCt method (fold expression=2^−ΔΔCt^). One-way ANOVA was used to determine statistical significance.

### Animals

Six-to-eight-week-old male C57BL/6J mice were used and housed in a specific-pathogen-free (SPF) facility at the Korea Advanced Institute of Science and Technology (KAIST). All procedures used in this study complied with guidelines and protocol (KA2021-004) of the Institutional Animal Care and Use Committee of KAIST.

### Virus infection and hIFN-λ3 variants treatment

For upper respiratory tract infection, mice were intranasally inoculated with 1 × 10^6^ plaque-forming units (PFU) of influenza A/PR/8/34 virus (H1N1, provided by HK Lee, KAIST, South Korea) in a 16μL volume under light anesthesia (4% isoflurane in oxygen). For intranasal hIFN-λ3s treatment, mice were intranasally administered 1 µg or 5 µg of hIFN-λ3-WT, hIFN-λ3-DE1 or G-hIFN-λ3-DE1 in 10 µL of PBS.

### Measurements of mouse ISG gene expression at nasal tissue by RT-qPCR

At 2 hours post hIFN-λ3 variant administration, nasal turbinates were isolated and homogenized in Trizol (#15596026, Invitrogen). Total RNA was extracted using the RNeasy Mini Kit (#74106, QIAGEN) according to the manufacturer’s protocol. cDNA was synthesized from 1 µg of RNA using an oligo-dT primer and the SuperScript III First-Strand Synthesis System for RT-PCR (#18080-051, Invitrogen). qPCR was performed using PowerUP SYBR Green Master Mix (#A25741, Applied Biosystems) on a QuantStudio 5 Real-Time PCR System (Applied Biosystems). Primer sequences are listed in Table S1 (Supporting Information). Ct values were normalized to Gapdh expression and relative gene expression was calculated using the ΔΔCt method (fold expression = 2^−ΔΔCt^).

### Immunofluorescence imaging

For immunofluorescence staining, mice were perfused with ice-cold PBS via the left ventricle to flush out blood, followed by fixation with 2% paraformaldehyde (PFA, #15710, EMS). For cryo-sectioning of the mouse nasal cavity, heads were post-fixed in 4% PFA overnight at 4°C, decalcified in 0.5 M EDTA (#ML005-01, Welgene) for 96 h at 4°C, and dehydrated by submersion in 30% sucrose (#SUC01, LPS) for 48 h at 4°C. Samples were embedded in Tissue-Tek® OCT™ compound (#4583, Sakura), frozen, and sectioned at 15 μm thickness. Frozen sections of nasal tissues were stained with 4′,6-diamidino-2-phenylindole (DAPI, #D-9542, Sigma-Aldrich) and mounted in Fluoromount-G (#0100-01, SouthernBiotech). Primary antibodies (1:300) included anti-influenza A (goat polyclonal, #ab20841, Abcam) and AF488-conjugated anti-mouse CD326 (Ep-CAM) (rat monoclonal, #118210, BioLegend). Secondary antibody (1:1000) was Alexa Fluor® 594 AffiniPure rabbit anti-goat IgG (H+L) (#305-585-045, Jackson ImmunoResearch). Images were acquired using a confocal microscope with 10× or 20× objectives (LSM980, Zeiss).

### Influenza virus titration

At 2 days post infection, nasal turbinate tissue was homogenized in minimum essential medium containing 0.3% BSA using a Precellys 24® bead-beating homogenizer (#P000669-PR240-A, Bertin Technologies) and lysing kit tubes (#P000911-LYSK0-A.O, Bertin Technologies). Homogenates were centrifuged (15 min, 2300 g) to clear debris, and viral titers in the supernatants were determined by standard plaque assay on Madin-Darby canine kidney cells (MDCK, provided by HK Lee, KAIST), as previously described[68].

## Data Availability Statement

The source data for all figures and supplementary figures are available as Source Data file. All the other data and materials used for the analysis are available from the corresponding author upon reasonable request. Source data are provided with this paper.

## Acknowledgements

We are grateful to the staff of the Research Solution Center at IBS. Computational work for this research was performed using the high-performance computing resources in the IBS Research Solution Center. This work was supported by the Institute for Basic Science (IBS-R030-C1 to H.M.K.), the National Research Foundation of Korea (RS-2025-00523615 to H.M.K. and RS-2023-00222762 to J.E.O.), and the KAIST Convergence Research Institute Operation Program.

## Contributions

J.Y. and H.M.K. designed the experiments and analyzed the data; J.Y. designed and purified the proteins, conducted protein stability tests, and performed biological activity experiments at the cellular level; S.Y. set up the nasal mucus mimic systems and conducted the protein diffusion assays; J.H.K. performed cross-reactivity and antiviral activity experiments in mouse models; L.F.V. wrote scripts for the computational protein design processes; J.Y., S.Y., J.H.K., L.F.V., M.C., H.J.C., J.E.O., and H.M.K. wrote the manuscript.

